# Beyond IC_50_ - A computational dynamic model of drug resistance in enzyme inhibition treatment

**DOI:** 10.1101/2024.03.21.586067

**Authors:** J. Roadnight Sheehan, Astrid de Wijn, Thales Souza Freire, Ran Friedman

## Abstract

Resistance to therapy is a major clinical obstacle to treatment of cancer and communicable diseases. Chronic myeloid leukaemia (CML) is a blood cancer that is treated with Abl1 inhibitors, and is often seen as a model for targeted therapy and drug resistance. Resistance to the first-line treatment occurs in approximately one in four patients. The most common cause of resistance is mutations in the Abl1 enzyme. Different mutant Abl1 enzymes show resistance to different Abl1 inhibitors and the mechanisms that lead to resistance for various mutation and inhibitor combinations are not fully known, making the selection of Abl1 inhibitors for treatment a difficult task. We developed a model based on information of catalysis, inhibition and pharmacokinetics, and applied it to study the effect of three Abl1 inhibitors on mutants of the Abl1 enzyme. From this model, we show that the relative decrease of product formation rate (defined in this work as “inhibitory reduction prowess”) is a better indicator of resistance than an examination of the size of the product formation rate or fold-IC_50_ values for the mutant. We also examine current ideas and practices that guide treatment choice and suggest a new parameter for selecting treatments that could increase the efficacy and thus have a positive impact on patient outcomes.

## 1 Introduction

Contemporary drug design relies on the identification of molecular targets, such as receptors or enzymes, that once activated or inhibited lead to a marked improvement in the disease state. One caveat of this paradigm is the risk for development of resistance where the disease to be treated is either cancer or a communicable (transmittable, pathogen borne) disease. Chronic myeloid leukaemia (CML) has been extensively studied as a model for the development of drug resistance especially when it comes to resistance due to mutations in the drug target.

CML is a blood cancer that in the vast majority of cases (95% [56]) involves a chromosomal translocation (the Philadelphia chromosome), which can be detected and is the basis for diagnosis. The chromosomal translocation results in a fusion of the breakpoint cluster region (BCR) of chromosome 22 with the Ableson leukemia (Abl) gene on chromosome 9. This BCR-Abl region encodes Abl1 that lacks its regulatory part. The deregulation promotes over-activity of the translated protein, Abl1 kinase, and this in turn disrupts cells’ processes, leading to: increased growth and multiplication; blocking of differentiation; and loss of programmed cell death [35]. The Philadelphia chromosome is not only the primary cause for CML, but also a contributing factor to other cancers [4].

Abl1 inhibitors, are the primary treatment for adults with CML [28]. The first Abl1 inhibitor treatment that is most commonly given to patients is imatinib, which has milder side effects than other Abl1 inhibitors [12]. Imatinib and other Abl1 inhibitors reduce cell proliferation, eliminating the symptoms of CML and rendering the presence of Philadelphia chromosomes undetectable [13]. However, treatment with Abl1 inhibitors does not provide a cure for the disease and patients often need treatment for the rest of their lives [42].

Although Abl1 inhibitor treatment is increasing survival rates, resistance to imatinib as initial treatment develops in around 25% of patients within two years [44] (estimations for other Abl1 inhibitors are: 20% for initial treatment with dasatinib within two years [15]; and 21% for secondary treatment after treatment of one prior tyrosine kinase inhibitor within one year [36]). In most cases, the resistance arises from mutations in the Abl kinase domain that alter the protein structure. If an Abl1 mutant is less affected by the Abl1 inhibitor, cells with this mutation then become prevalent in the cell population, and increase the growth rate [41]. Table 1 gives information about the Abl1 inhibitors and the mutations that are associated with resistance to those treatments that we examine in this work.

**Table 1:** The table (adapted from [10]) shows the mutants associated with resistance for imatinib, ponatinib, and dasatinib. These mutants were considered in this work as they are the most common and have been extensively studied. * Although these mutations do have an increased IC_50_ for ponatinib, they do not lead to resistance alone, but do so as compound mutations.

| Drug | imatinib | ponatinib | dasatinib |
| --- | --- | --- | --- |
| **Mutants considered in this work** | G250E<br>E255K<br>E255V<br>T315I<br>T315M<br>Y253H | E255K*<br>E255V*<br>T315I* | T315I<br>T315M |

Once significant cell populations with imatinib resistant mutations arise, the treatment must change. Second- and third-generation Abl1 inhibitor treatments are available. However, which drug should be matched to which mutation is not always clear, and, overall, there is a need to develop treatment protocols that will minimise the risk for resistance [43], which requires an understanding of resistance mechanisms that take into account the effects of drug binding, enzyme activation [22, 20] and interference with catalysis [26].

In this paper, we develop a computational model to assess whether the product formation rate of the Abl1 enzyme and its mutations provides an indicator of resistance to Abl1 inhibitor treatment. We thereafter compare the leading approach of drug selection in CML treatment (relative IC_50_, see section 2) with methods that include further information (such as Abl1 inhibitor concentration data) to assess their ability to infer drug resistance, which would better inform choices in treatment when resistant mutations arise.

## 2 Approach to understanding resistance

Many factors have been examined in the pursuit of finding the best choice of Abl1 inhibitor to switch to once resistant mutations arise. Often the data of the wildtype (WT) Abl1 enzyme is compared to that of the mutation (Mut) - specifically, the IC_50_ values. IC_50_ is the concentration of the inhibitor in which the enzyme’s activity or cell’s growth (depending on the assay) reduces to half of its maximum value. Typically in drug selection, an examination of the relative IC_50_ of each Abl1 mutation is used: a ratio of 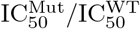. This is often described as an imperfect approach [60, 23, 32]. Generally, the lower the value of the IC_50_, the greater the effect of the inhibitor. However, 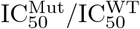 quantifies which mutations a drug is most effective against, rather than which drug is best suited to treating each mutation. Furthermore, there are other factors that are relevant for choosing the treatments. A patient’s tolerance and the safety profile of the treatments must be considered as well.

An additional complication for the selection process is that the mutations that give rise to the mutant Abl1 enzymes affect other processes in the cell. For example, the catalytic rate constant, *k*_cat_, varies between mutations, indicating that different mutations may provide a higher product formation rate. However, due to other unknown effects on other processes in the cell, it is unwise to assume that the size of *k*_cat_ is solely responsible for (or has any effect on) whether a mutation is resistant to a certain treatment.

Another method of examining and comparing Abl1 with its mutations is catalytic efficiency:

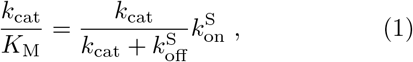

where *K*_M_ is the Michaelis constant for the enzyme; 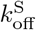 is the rate constant of the substrate unbinding without catalysis from the enzyme; and 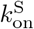 is the rate constant of the substrate binding to the enzyme per unit concentration of the substrate. A compound mutation is a double mutation in the same allele (rather than one mutation in one allele and another mutation in another allele). In cases where two mutations do not show resistance alone, but do so as a compound mutation, the compound mutation might have higher catalytic efficiency than either of the single mutations [26]. This highlights the possibility that the catalytic efficiency might affect the resistance, and therefore we consider the catalytic efficiency in our investigations.

In this paper, we take a mathematical, computational approach to investigate the change in the product formation rate of the wild-type (WT) BCR-Abl Abl1 enzyme and six of its mutations when subjected to treatment with one of three different Abl1 inhibitors. Our aim is to provide insight into how these changes in the product formation rate lead to drug resistance. We also investigate analytically the link between a drug and mutation combination showing resistance and different measurable quantities of the mutant Abl1 enzymes (such as *k*_cat_, *K*_M_ and IC_50_).

The Abl1 inhibitors in focus in the work are imatinib, ponatinib and dasatinib. With the computational model, we examine these inhibitors in combination with the WT Abl1 enzyme and six mutations of it: G250E, E255K, E255V, T315I, T315M, and the compound mutation Y253H-E255V. When analytically investigating different possible indicators of resistance, we also examine the mutations Y253H, G250E-T315I, Y253H-T315I, and E255V-T315I.

## 3 Methods

The success of Abl1 inhibitor in the clinic shows that enzyme inhibition is coupled to a reduction in the product formation rate of Abl1. In a clinical setting, treatment is monitored through hematologic (peripheral blood counts of leukocytes, platelets, and immature cells), cytogenetic (the prevalence of the Philadelphia chromosome in cells), and other molecular measures of blood or bone marrow samples from patients [58]. These tests are population counts and considering the Abl1 enzyme is a key part of growth and multiplication signalling, this might suggest that the product rate is a good indicator of resistance. Consequently, we use computational modelling to describe the product rate for different drug resistant mutations under different treatments. Based on our model, we analytically derive an indicator of the severity of the resistance in drug-mutation combinations that can be used in practical settings to better inform treatment choice.

### 3.1 Modelling of drug resistant mutations

To establish a dynamic model, we consider the different states that a system consisting of an enzyme, substrate, and inhibitor can be in and derive a complete set of time-dependent rate equations for the concentrations of enzyme in each state,

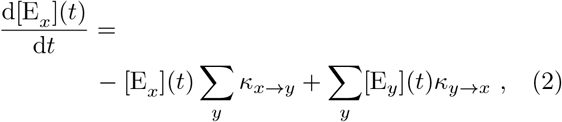

where *x* and *y* are the labels of the enzyme states. We denote the concentration of the enzyme in state E_*x*_ by [E_*x*_], and *κ*_*x*→*y*_ is the rate constant for transition from state E_*x*_ to state E_*y*_.

To set up this model, we need two components: the states that need to be considered, and the transition rates between them. In section 3.1.1, we determine the relevant states of the system. Thereafter we examine the system by assuming quasi-equilibrium conditions in section 3.1.2. Based on this, we then derive the relationships between the rate constants. Quantitative values and approximations for the rate constants can then be made based on existing experimental results and simulations, which we describe in section 3.1.3.

To more accurately model the physiology of drug treatment, where the concentration fluctuates, we also consider the pharmacokinetics of the inhibitor in the system as described in section 3.1.4. In section 3.1.5, we outline the calculations of the initial conditions of the system analytically, assuming a pre-existing quasiequilibrium steady state before treatment.

#### 3.1.1. Determining the states of the system

As the Abl1 enzyme is part of a signalling pathway that controls cellular growth, its deregulation results in rampant cell proliferation. The cellular system is complex, but as the Abl1 signalling is of high importance in the pathway, the system can be simplified to focus on one element of CML in patients - the phosphorylation of tyrosine residues in various proteins by Abl1, i.e. the product formation rate. The importance of this mechanism in CML is verified by its successful treatment with Abl1 inhibitors [59]. Focusing on the product formation rate as a measure for the efficacy of the drugs allows us to quantify and compare the many drug-resistant Abl1 enzymes with small changes in the BCR-Abl gene.

We first examine the possible states that the system can be in. The enzyme can be active or inactive. Moreover, it can be bound or not to the inhibitor, the substrate protein, and ATP. There are many combinations of these that could be considered. However, we can eliminate some of these combinations from our simplified computational model. Firstly, the site that the inhibitory drugs bind to is the same site that the ATP molecule binds to, therefore the ATP and an inhibitor cannot be bound to an Ab1 enzyme at the same time. Furthermore, if bound to an inhibitor drug molecule, the flexibility of the protein structure of the enzyme is reduced and prevents the binding of the substrate protein to Abl1 [50]. Thus, a combination of the inhibitor and the substrate protein binding with the enzyme together does not need to be considered in our model. The likelihood of the substrate binding to the inactive form of the Abl1 is very small [49]; we therefore do not consider the states with binding of the substrate to the inactive enzyme in our model. We also combine multiple transitions into single transitions between states (i.e. active unbound state to active bound to both the substrate protein and ATP, without transition via binding to one of the molecules). This leaves a total of five different states for the system: active enzyme not bound to anything; inactive enzyme not bound to anything; active enzyme bound to both the substrate protein and an ATP molecule; active enzyme bound to an inhibitor; and inactive enzyme bound to an inhibitor, as shown in Figure 1. Figure 1 also shows the transition rate constants between states, which are discussed in detail in subsection 3.1.3.

**Figure 1:**
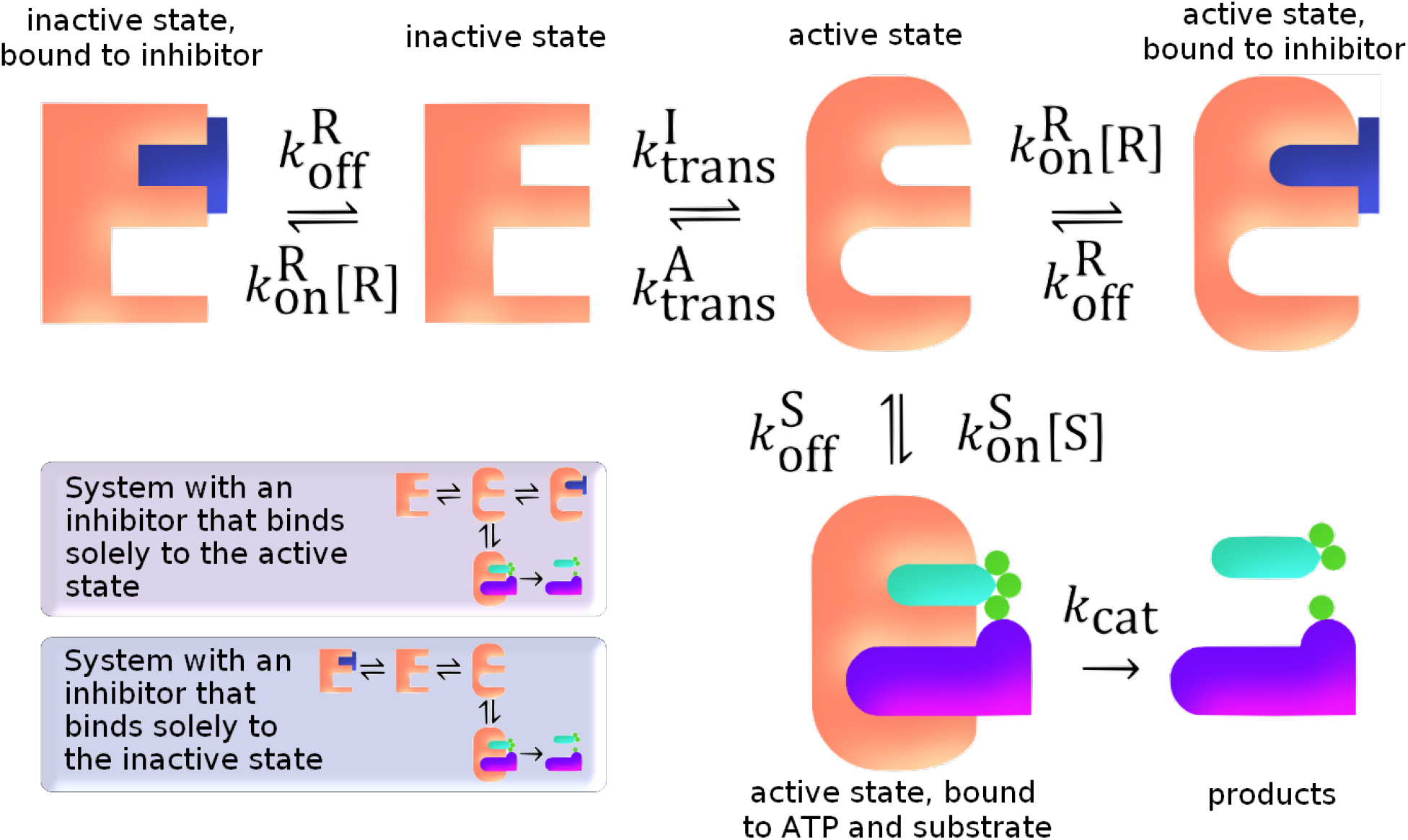
A simplified system of the Abl1 enzyme to be modelled. The states, molecules they bind to, the transitions between states, and the rate constants for each transition are shown. The Abl1 enzyme is represented by the peach-coloured E-shape in both its active (round corners) and inactive (sharp corners) states; the inhibitor is illustrated by the dark blue T-shape; and the remaining shapes depict the ATP with its phosphate groups and the protein substrate that is phosphorylated. Transition arrows are accompanied by their rate constants. Two smaller diagrams in the bottom left corner show the states of the system in the cases of active-only binding and inactive-only binding inhibitors.

Finally, we can reduce the number of possible states further by considering the binding of the different inhibitor drug molecules to the enzyme. Imatinib and ponatinib both bind only to the inactive enzyme state while dasatinib binds more strongly to the active enzyme state. This leaves just four states of the Abl1 in each simulation.

#### 3.1.2 Fixed conditions and quasi-equilibrium

In order to develop our model for the system dynamics with fluctuating inhibitor concentration, it is useful to first consider the system with fixed inhibitor concentrations and assume a quasi-equilibrium. This system can be used to relate the various rate constants to one another, and can serve as a basis to construct equations for the dynamics.

Under fixed conditions, the system behaves in a relatively straight-forward way that can be determined using experimental and simulation data. We denote the concentration of substrate with [S] and the concentration of inhibitor with [R]. We assume that ATP is in surplus relative to the substrate and its concentration is therefore not treated explicitly by the model. The free-energy difference between the active and inactive enzymes states is denoted as Δ*G*^I→A^, which is defined as *ϵ*_I_ − *ϵ*_A_ where *ϵ*_I,A_ are the free energies of the inactive and active state of the enzyme, respectively. Using principles of chemical kinetics [19] and statistical mechanics, we can find the proportions of the enzyme concentration that correspond to each of the states in quasi-equilibrium described in Figure 1. These proportions of the enzyme concentration are referred to as relative weights and are shown in Table 2.

**Table 2:**
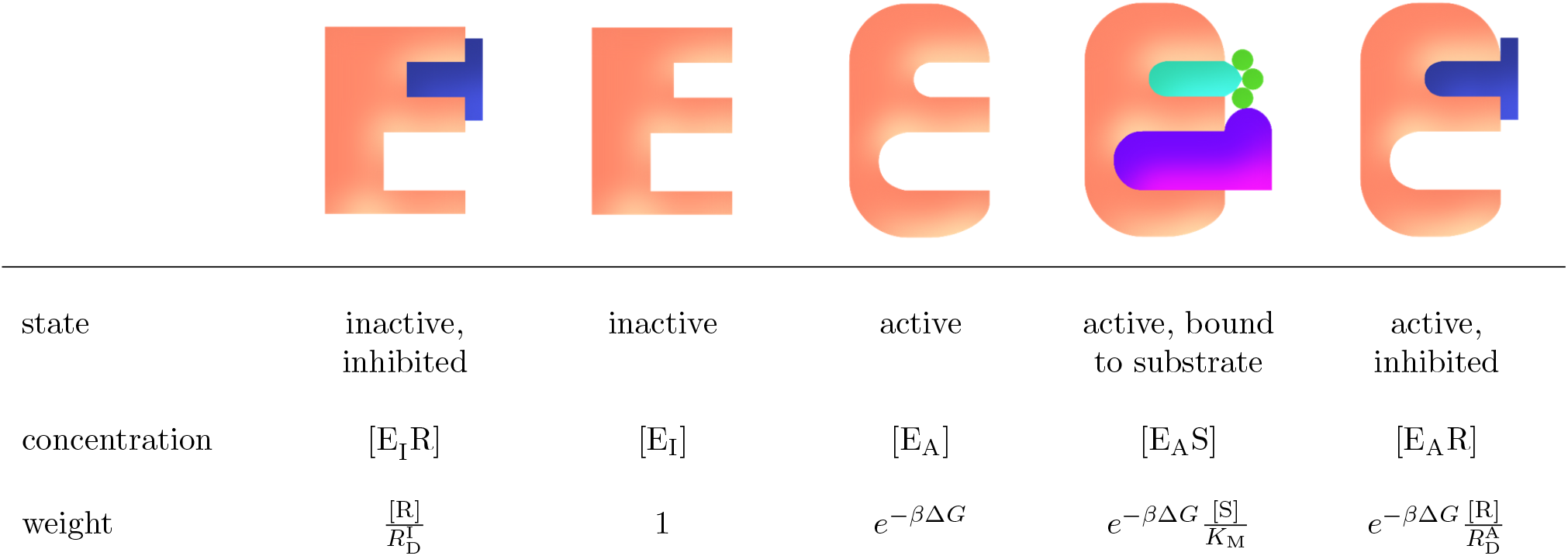
The enzyme states and their relative weights within the system in quasi-equilibrium. [R] is the concentration of the inhibitory drug in the system; 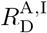 is the dissociation constant of the binding of the inhibitory drug the the enzyme in either active (A) or inactive (I) state; *β* is the reciprocal of the product of Boltzmann’s constant and the temperature (*β* = 1*/k*_*B*_*T*); Δ*G* is the change in Gibbs free energy between the active and inactive enzymes; [S] is the concentration of the substrates (it is assumed that ATP is in surplus relative to the substrate and its concentration is therefore not treated explicitly by the model); and *K*_M_ is the Michaelis constant of the binding of the substrate to the active enzyme state.

We can now express the product formation rate, 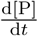, for this system in terms of the concentration of active enzyme bound to the substrate [E_A_S], as in [19]:

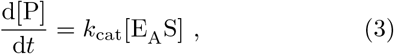

where *k*_cat_ is the turnover number (also known as the catalytic rate constant) for the active enzyme. The concentration of active enzyme bound to the substrate can be expressed as a proportion of the total concentration of enzymes, [E_tot_], in quasi-equilibrium. This is done using the weight for the state [E_A_S] from Table 2 as a fraction of the total if the weightings for the system (which differs depending on the inhibitor present). Thus giving the product formation rate as:

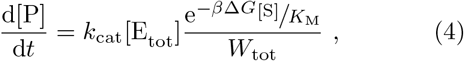

where the total weight, *W*_tot_, is different depending on whether an inhibitor is present, and which inhibitor that is. For imatinib (i) and ponatinib (p) which bind to the inactive state of the enzyme (see Figure 1) the total weight is:

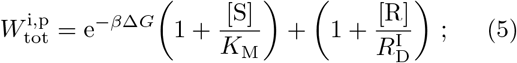

and for dasatinib (d) which binds to the active state of the enzyme (see Figure 1) the total weight is:

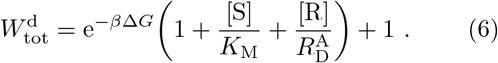

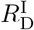 and 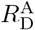 are the inhibitor dissociation constants.

#### 3.1.3 Quantitative determination of rate constants

The real inhibitor concentration in patients varies with time, which in turn leads to fluctuations in the product formation rate, and potentially, a shift in the average. A more detailed discussion on and calculation of the fluctuations in the inhibitor concentration can be found in section 3.1.4. To take this into account, we must model the time-dependence of the concentrations of all the states and thus describe not only the quasi-equilibrium relative weights, but also the transition rates between states. These are modelled by the rate constants (*κ*_*x*→*y*_ and *κ*_*y*→*x*_) defined in Equation 2. In our model we calculate the rate constants as depicted in Figure 1 and expanded in Table 3. Our approach to obtaining these rate constants is described below.

**Table 3:** To bridge two naming conventions (one more suitable to mathematical notation and one more conventional in enzymology), the table matches the rate constants as described in Equation 2 to those labelled in Figure 1.

| rate constant<br>of transitions<br>as in Equation 2 | rate constant<br>as labelled in<br>Figure 1 |
| --- | --- |
| $\kappa_{[E_I R] \rightarrow [E_I]}$ | $= k_{\text{off}}^R$ |
| $\kappa_{[E_I] \rightarrow [E_I R]}$ | $= k_{\text{on}}^R [R]$ |
| $\kappa_{[E_I] \rightarrow [E_A]}$ | $= k_{\text{trans}}^I$ |
| $\kappa_{[E_A] \rightarrow [E_I]}$ | $= k_{\text{trans}}^A$ |
| $\kappa_{[E_A] \rightarrow [E_A R]}$ | $= k_{\text{on}}^R [R]$ |
| $\kappa_{[E_A R] \rightarrow [E_A]}$ | $= k_{\text{off}}^R$ |
| $\kappa_{[E_A] \rightarrow [E_A S]}$ | $= k_{\text{on}}^S [S]$ |
| $\kappa_{[E_A S] \rightarrow [E_A]}$ | $= k_{\text{off}}^S$ |
| $\kappa_{[E_A S] \rightarrow [E_A] + \text{Products}}$ | $= k_{\text{cat}}$ |

##### Abl1 inhibitor binding rate constants: 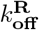 and 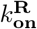

To obtain 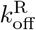 we can make use of the experimentally determined inhibitor residence times (*t*_R_) from [38]. The residence time is a measure of the time the molecule is bound until its unbinding. The number of molecules unbinding from a bound state that occur per unit of time gives an unbinding rate, i.e. 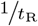. From this we can obtain the 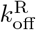 for each Abl1 inhibitor for the WT:

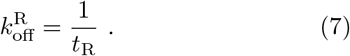

These values can be found in Table 4.

**Table 4:** Residence times, *t*_R_, of the inhibitor molecules unbinding from the wild-type from [38] and the rates of 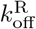 calculated from them.

| ABL1<br>inhibitor | $t_R$<br>(min) | $k_{\text{off}}^R$<br>(min<sup>-1</sup>) |
| --- | --- | --- |
| imatinib | 17 | 0.059 |
| ponatinib | 205 | 0.00488 |
| dasatinib | 249 | 0.00233 |

For the rate constant in the other direction, 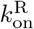, we will make use of the quasi-equilibrium between the two rates, through the general definition of the dissociation constant of the inhibitor given as:

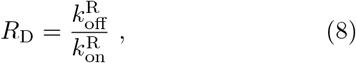

where 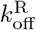 is the rate constant of the inhibitor unbinding from the enzyme (measured in min^−1^) and 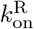 is the rate constant of the inhibitor binding per unit concentration of the inhibitor (usually measured in *µ*M^−1^min^−1^).

To obtain the dissociation constant, we use an approach similar to [19], making use of the experimentally measurable IC_50_ value of the Abl1 inhibitor. We compare the product formation rate in Equation 4 for [R]= 0 and [R]=IC_50_. As the IC_50_ is defined as the concentration of inhibitor needed to halve the product formation rate, we can write:

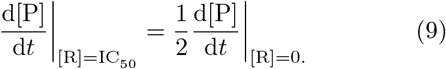

Substituting [R]=IC_50_ into Equations 5 and 6 and re-evaluating Equation 4, yields expressions for the inhibitor dissociation constants for the active-binding case, 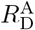, and the inactive-binding case, 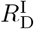:

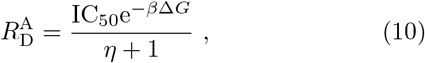

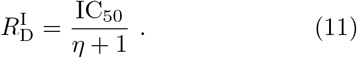

Here *η* is used as short hand for

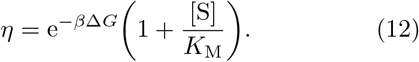

The IC_50_ is experimentally known [55, 64]. We obtain a value for Δ*G* from molecular simulations, which are discussed in detail later when examining the rate of transition between the active and inactive states. The total substrate concentration [S] is chosen to be constant (i.e. an assumed perfect replenishment rate) and estimated to be 10 *µ*M. Table 5 shows the *K*_M_ values used in this work, obtained from [26], and the IC_50_ values, from [55] and [64].

**Table 5:**
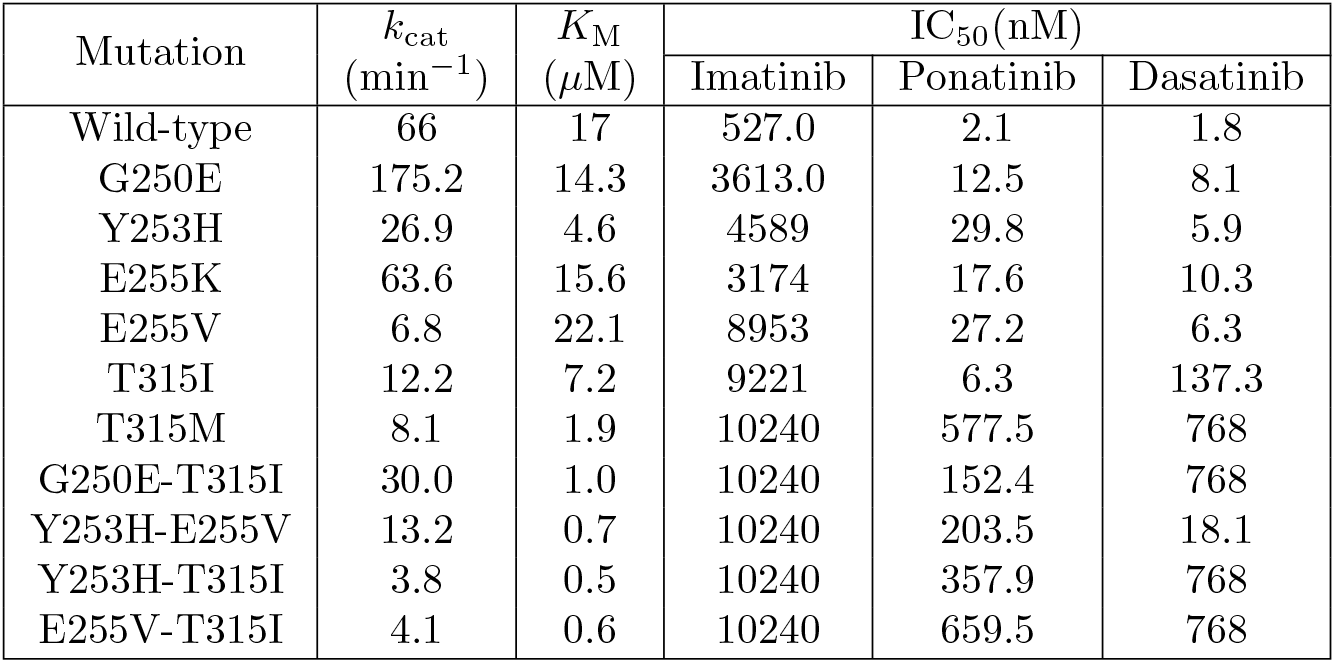
The values for *k*_cat_ and *K*_M_ are sourced from [26]. All the IC_50_ values for the wild type, G250E, E255K, E255V and T315I are from [55]. The IC_50_ for imatinib for Y253H is also from this source. The other IC_50_ values for Y253H along with all IC_50_ values for T315M, G250E-T315I, Y253H-E255V, Y253H-T315I, and E255V-T315I are from [64].

| Mutation | $k_{\text{cat}}$<br>($\text{min}^{-1}$) | $K_{\text{M}}$<br>($\mu\text{M}$) | $\text{IC}_{50}(\text{nM})$ | | |
| --- | --- | --- | --- | --- | --- |
|  |  |  | Imatinib | Ponatinib | Dasatinib |
| Wild-type | 66 | 17 | 527.0 | 2.1 | 1.8 |
| G250E | 175.2 | 14.3 | 3613.0 | 12.5 | 8.1 |
| Y253H | 26.9 | 4.6 | 4589 | 29.8 | 5.9 |
| E255K | 63.6 | 15.6 | 3174 | 17.6 | 10.3 |
| E255V | 6.8 | 22.1 | 8953 | 27.2 | 6.3 |
| T315I | 12.2 | 7.2 | 9221 | 6.3 | 137.3 |
| T315M | 8.1 | 1.9 | 10240 | 577.5 | 768 |
| G250E-T315I | 30.0 | 1.0 | 10240 | 152.4 | 768 |
| Y253H-E255V | 13.2 | 0.7 | 10240 | 203.5 | 18.1 |
| Y253H-T315I | 3.8 | 0.5 | 10240 | 357.9 | 768 |
| E255V-T315I | 4.1 | 0.6 | 10240 | 659.5 | 768 |

We can use these 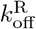 values for the WT; Equations 10 and 11 for the dissociation constants; and Equation 8 to calculate the 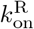 values for the WT. These inhibitor associated rate constants for the WT can be used to estimate the rate constants of the mutants based on what we know of the working mechanisms of the inhibitors. We assume that for the inactive-binding Abl1 inhibitors (imatinib and ponatinib), the 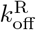 remains the same across the WT and the mutants. This is based on a study that has shown that, once bound, a drug interacted with resistance associated mutants of a kinase in a similar manner to its interaction with the wild type enzyme [21]. Similarly, we can assume the 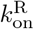 is the same across the WT and all mutations for active-binding Abl1 inhibitors (dasatinib). The resulting values for the dissociation constants as well as for the binding and unbinding rate constants for the Abl1 inhibitors can be seen in Table 6.

**Table 6:**
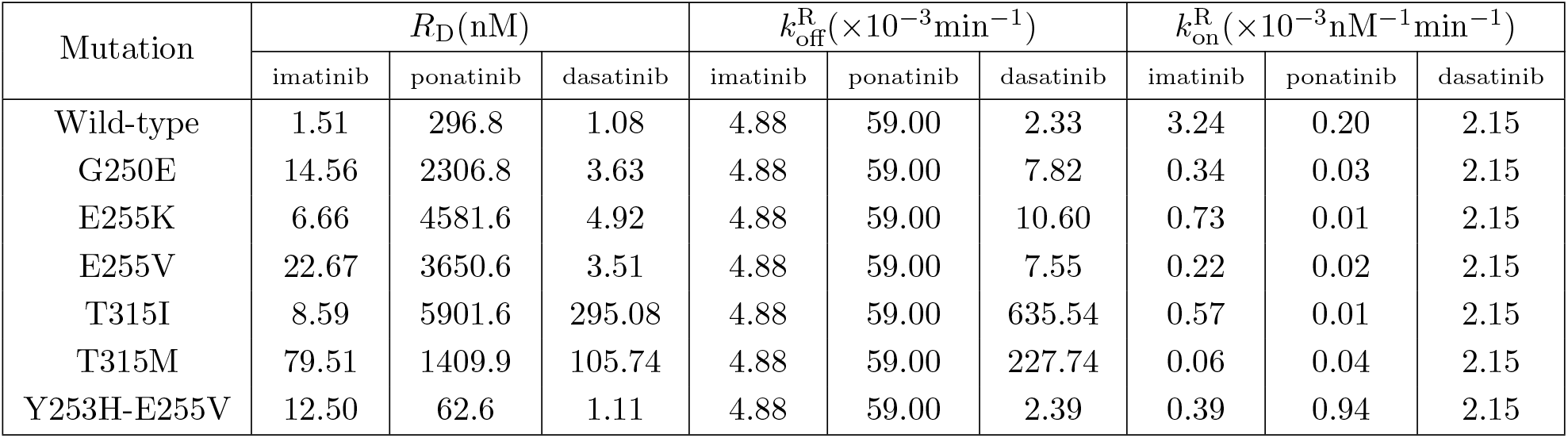
The values calculated for the inhibitor dissociation constant, as detailed in Equations 11 and 10 and approximated values for the binding rate constants of the inhibitors.

##### Substrate binding: 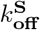 and 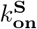

The rate constants for binding and unbinding of the substrate to the enzyme can be estimated from experimental results of the product formation rate, in this case the coefficients *k*_cat_ and *K*_M_ for the WT and each mutation. As we know most about the behaviour of the WT, we will use this as a starting point to compare the behaviour of the mutants to. In the following, we use superscript to indicate the WT or mutant (Mut). For the WT, 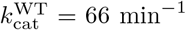 [26]. With the WT we expect the enzyme to be more likely to catalyse than unbind, so that 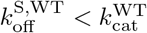, while the rate constants should still be of the same order of magnitude. We therefore estimate 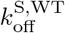 by 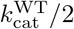, i.e. 33 min^−1^. Using the relationship:

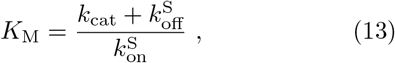

we can calculate a value for 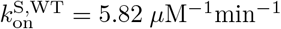.

For the mutants, which in the absence of inhibitor are less fit than the WT, we expect the opposite, that the enzyme is more likely to unbind than to catalyze - i.e. 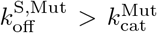, while the rate constants should again be of the same order of magnitude. We therefore use as a first estimate 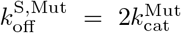, which means that 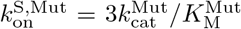. However, the mutants should also not be so unfit as to not be able to compete at all with the WT when there is no treatment, meaning that 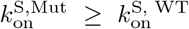. If this condition is not met, then we increase both 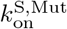 and 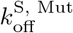 until it is. In this case 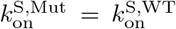 and 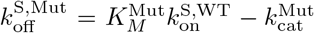. The final estimates are shown in Table 7.

**Table 7:** The values approximated and calculated for 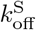 and 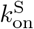 for the WT and each mutant Abl1 enzyme.

| Mutation | $k_{\text{off}}^S$<br>($\text{min}^{-1}$) | $k_{\text{on}}^S$<br>($\mu\text{M}^{-1}\text{min}^{-1}$) |
| --- | --- | --- |
| Wild-type | 33.0 | 5.82 |
| G250E | 350 | 36.8 |
| E255K | 127 | 12.2 |
| E255V | 122 | 5.82 |
| T315I | 29.7 | 5.82 |
| T315M | 16.2 | 12.8 |
| Y253H-E255V | 26.4 | 56.6 |

##### Transitions between the active and inactive enzyme states: 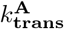 and 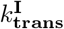

The ratio of the transition rate constants between the active and inactive enzymes is related to the freeenergy difference Δ*G*^I→A^ through:

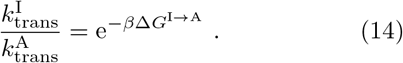

For the mathematical, computational model this needs to be found as it is not directly measurable experimentally. Using free energy perturbation (FEP) simulations, we are able to determine the change in Δ*G* between each mutation and that of the WT:

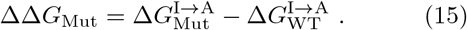

To find the structure of the active and inactive Abl1 WT enzyme, the X-ray crystal structures from the *Protein Data Bank* (PDB) of the Abl1 WT enzyme bound to a dasatinib molecule (PDB ID 2GQG) [63] and bound to an imatinib molecule (PDB ID 2HYY) [16] were superimposed by sequence in *BIOVIA Discovery Studio* version 2021 [9]. As dasatinib only binds to the active state of the Abl1 enzyme, this was used to find the active state. Similarly, imatinib is an inactive-binding inhibitor, so this provided the inactive state. The inactive structure had the smaller sequence and one residue missing in the middle of the chain. Due to this, the extra residues on the active structure were deleted and the missing residue was added to the inactive structure. The active state was phosphorylated in the crystal structure. To make the atomistic composition the same, the phosphate group was removed. After these changes by *BIOVIA Discovery Studio*, both models were corrected by *Swiss-Model* online tool [62].

Molecular dynamics (MD) simulations were thereafter carried out using *GROMACS* version 2021.5 [8, 61, 1]. The *Amber99sbmut* force field (modified version of *Amber99sb* used by *GROMACS*’ plugin *pmx* to handle mutations [24, 25]) was used for the solute atoms, and the transferable intermolecular potential with 3 points model (TIP3P) was employed for water molecules [34]. A cutoff distance of 1.2 nm was used to compute van der Waals and Coulomb interactions. Long-range electrostatic interactions were modelled using the Particle Mesh Ewald (PME) method [18]. The LINCS algorithm [31, 30] was used to constrain bonds involving hydrogen atoms in the solute, and the SETTLE algorithm [46] was employed to the same aim with water molecules. The simulations were performed at constant temperature of 300 K, using the velocity rescaling thermostat [11] (*τ*_T_ = 0.1 ps), and at constant pressure of 1 bar. The pressure was kept constant by the Berendsen barostat [7] for the equilibration and by the Parrinello-Rahman barostat [51] for the transitions runs (with *τ*_P_ = 1 ps).

The *pmx* package was used for preparation of the topologies [57, 25]. Mutations (E255K, E255V, G250E, T315I, T315M and Y253H-E255V) were generated with the `pmx mutate` module. Hybrid topologies, containing information about the WT and the mutated state of the enzyme, were built from these PDB files, with the `pmx gentop` module. The system was built as a dodecahedron box, placing the enzyme at the center, at a minimal distance of 1.2 nm from each edge of the box. The protein was solvated, adding K^+^ and Cl^−^ ions to neutralize the charges and reach the concentration of 0.15 mM. For each mutation, a single energy minimisation was performed using a steepest descent algorithm followed by a short MD simulation of 20 ps, where positional restraints were imposed on all solute heavy atoms in order to equilibrate the water and ions around the protein. Next, the system was equilibrated for 10 ns after removing the restraints. The equilibration was repeated ten times for each state of the enzyme to ensure better accuracy through sampling. One hundred snapshots were extracted from the last 8 ns of the equilibration trajectories. From each of these snapshots, a single short MD simulation of 100 ps was spawned (200 ps for Y253H-E255V), starting from the WT (forward transition) and from the mutant (reverse transition). This resulted in simulations of 100 transitions in each direction. The Parrinello-Rahman barostat [51] was used for these simulations, and velocities were generated at each transition run. The *GROMACS* command `gmx mdrun -dhdl` was used to calculate the change in the Hamiltonian for each simulation step, where the integral along the whole path corresponds to the work done in that specific run. Finally, the average Gibbs energy for each mutation was obtained with Bennett Acceptance Ratio [6] using the `pmx analyze` module over all values and averaging over the ten simulations. The above setup was applied for both active and inactive enzymes.

From the FEP simulations we can obtain the difference in Gibbs free energy upon mutation between active and inactive states 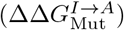. The same information could be obtained performing two transition (inactive to active) simulations, one with the WT native protein and another after mutation. This can be seen on the closed thermodynamic cycle (equation 16).

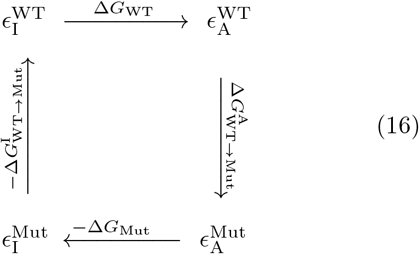

As the clockwise sum in Equation 16 is equal to zero, we can write the following equations:

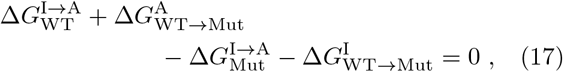

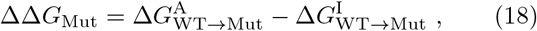

and

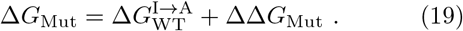

The values for 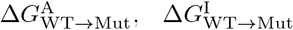, and ΔΔ*G*_Mut_ can be found in Table 8. It is assumed that the WT enzyme would prefer active state over inactive state, i.e. 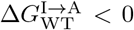. The size of 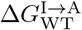 was estimated as −1 kcal/mol [49].

**Table 8:** The values for 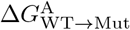 and 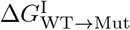 with the resultant ΔΔ*G*_Mut_, found through FEP simulations. ΔΔ*G*_Mut_ for the wild-type is 0 kcal/mol.

| Mutation | $\Delta G_{\text{WT} \rightarrow \text{Mut}}^{\text{A}}$<br>(kcal/mol) | $\Delta G_{\text{WT} \rightarrow \text{Mut}}^{\text{I}}$<br>(kcal/mol) | $\Delta \Delta G_{\text{Mut}}$<br>(kcal/mol) |
| --- | --- | --- | --- |
| G250E | -60.45 ± 0.6 | -62.66 ± 0.1 | -2.2 ± 0.6 |
| E255K | 65.60 ± 0.3 | 65.11 ± 0.5 | -0.5 ± 0.6 |
| E255V | 57.12 ± 0.2 | 56.77 ± 0.2 | -0.4 ± 0.3 |
| T315I | 29.72 ± 0.1 | 29.82 ± 0.2 | 0.1 ± 0.2 |
| T315M | 28.40 ± 0.3 | 28.50 ± 0.2 | 0.1 ± 0.3 |
| Y253H-E255V | 56.66 ± 0.4 | 57.06 ± 0.3 | 0.4 ± 0.4 |

In order for the imatinib to be an effective drug against the WT, the inactive state must be fairly accessible. On this basis, and given 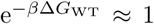, we have assigned that 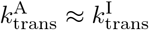 for the WT and they are of the order of 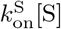. This then provides the foundation for the remaining values of 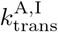 for the mutations. As the values of ΔΔ*G*_Mut_ for most of the mutations are small, with the exception of G250E, we can assume that the values for 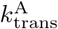 and 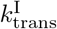 for these mutations are similar to those of the WT. The sizes of these rate constants for G250E will be assumed to be slightly smaller due to its larger ΔΔ*G*_Mut_. With all this information in mind, we have approximated that 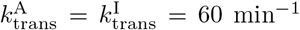 for all mutants, with the exception of G250E in which 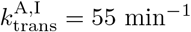.

#### 3.1.4 Time-dependent drug concentrations in patients

In reality, the drug concentration in the plasma of a patient is not constant, but fluctuates based on the timing between drug intake and physiological conditions. Therefore, for the time-dependent drug concentration in patients, we assume a perfect ‘model’ patient who receives a regular daily oral dosage of the drugs in a course of treatment with a fixed time between doses and whose body consistently maintains other biological conditions so that absorption and elimination of the drug happen in the same way every single day.

After the first dose of treatment, the concentration of the drug in the patient increases. There is a sharp absorption curve as the drug is absorbed into the bloodstream. The body begins to eliminate the drug as soon as it enters the system, however, the elimination starts off very small. At a point, the process of the drug being eliminated from the system becomes dominant and the concentration falls exponentially, usually with a long tail. The concentration continues to drop, but is still present at the point in which another dose of treatment is to be taken. With the second dose of treatment, there will be another sharp absorption curve, but this time the concentration it reaches is higher, as there is still some of the first dose remaining in the system. The remainder from the dose when the next dose is taken contributes to a taller peak in concentration until a steady-state of concentration is reached where the dosing creates a fluctuating concentration between a consistent daily minimum and maximum.

We solve for the plasma concentration of the inhibitor *C*(*t*) at time *t* since the first dose of Abl1 inhibitor. Let *n* be the number of days (24 hour periods) since the first dose. *k*_*a*_ and *k*_*e*_ are the absorption and elimination rate constants of the Abl1 inhibitor. The solution for the multiple dose pharmacokinetics for the plasma concentration of the inhibitor *C*(*t*) is given by [48]

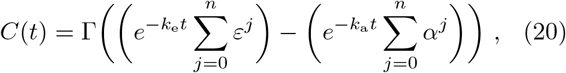

where Γ, *α*, and *ε* are constants that are given by:

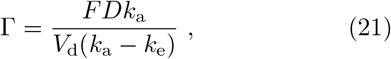

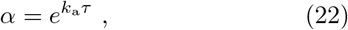

and

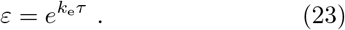

*F* is the bioavailability, *D* is the daily dose taken in moles, *V*_*d*_ is the volume of distribution, and *τ* is the dosing interval. The bioavailability of imatinib is 98% [52], but only qualitative descriptions for the bioavailability of ponatinib and dasatinib were found in the literature (“high” [2] and “potent, orally bioavailable” [40], respectively). Therefore, the value for *F* was assumed to be 1 across all three drugs. The elimination rate constant, *k*_e_, was found using half-life values, 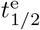, as: 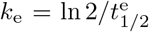. Medicinal doses, *D*, are provided as a mass of the active ingredient - the molar mass was used to calculate the dose in moles. We chose a system with daily doses of treatment, hence the time between doses, *τ*, is 24 hours. For the other values that shape the inhibitor concentration dynamics for the three drugs of interest, see Table 9.

**Table 9:** Values for the calculation of the time-dependent drug concentration in the system based on observed data found and derived from a variety of sources. Absorption constants, *k*_a_ where taken from [27, 29], and [17]. The elimination half-lifes, 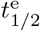, that the elimination constants, *k*_e_, were calculated from are from [53, 33], and [37]. For the dose, *D*, we chose masses described as daily doses in”typical treatment” [14, 54], and [39]. The volumes of distribution, *V*_d_, were taken from [53, 45], and [37]. Values for Γ, *α* and *ε* were calculated using Equations 21-23.

|  | Inhibitor drug |  |  |
| --- | --- | --- | --- |
|  | imatinib | ponatinib | dasatinib |
| $k_a$ (h<sup>-1</sup>) | 0.94 | 1.302 | 1.74 |
| $t_{1/2}^e$ (h) | 18 | 24 | 4 |
| $k_e$ (h<sup>-1</sup>) | 0.0385 | 0.0289 | 0.173 |
| $D$ (mg) | 400 | 45 | 180 |
| $M$ (g/mol) | 493.603 | 532.6 | 488.01 |
| $D$ (μmol) | 810.4 | 84.49 | 368.8 |
| $V_d$ (L) | 435 | 1223 | 2502 |
| $\Gamma$ (nM) | 2112.46 | 70.65 | 163.73 |
| $\alpha$ | 6.28E+09 | 3.72E+13 | 1.37E+18 |
| $\varepsilon$ | 2.52 | 2 | 64 |

#### 3.1.5 Statistical mechanics of the initial conditions of the system without treatment

The initial state from which the simulations start should be equivalent to the untreated condition. We use an analytical approach to determine this state. As the initial conditions are a system with no Abl1 inhibitor present, the system should reach a quasi-equilibrium steady-state. Using the results from section 3.1.2 and removing the inhibited states, we can write the steady-state distribution of a system without inhibitor as:

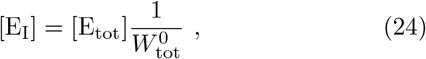

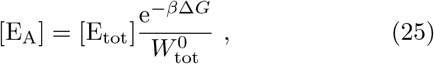

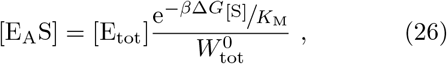

where,

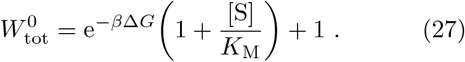

We set the total concentration of the enzyme, [E_tot_], to 1 *µ*M [5, 47].

### 3.2 Analytical approach to Abl1 inhibitor selection in treatment to combat drug resistance

To determine the effectiveness of treatment options on a specific mutation, in this work, we base ourselves directly on a comparison between the resulting product formation rates in our model. The maximum product formation rate, or velocity, for a given enzyme concentration is denoted by *V*_max_, which is proportional to *k*_cat_. The product formation rate of a system can be described in terms of *V*_max_, *K*_M_, and the substrate concentration, [S] [3].

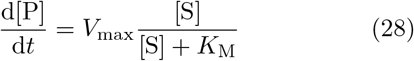

Considering just the enzyme and substrate (not treatment or other external factors), there are two main routes by which a mutation can affect this expression, either through *V*_max_ via *k*_cat_, or through *K*_M_ as the fraction [S]*/*([S] + *K*_M_). As mentioned in section 2, catalytic efficiency is also a candidate for the prediction of treatment resistance. These will be presented and examined in section 4.2.

A similar equation can be derived for the product formation rate when [R]= 0 and [R]= IC_50_ to derive an equation for how the product formation rate is affected by the inhibitor concentration, [R]. For this we can write the product formation rate for non-zero values of [R] in terms of the product formation rate with no inhibitor present ([R]= 0). We fist consider Equation 4 and note that the inhibitor concentration only enters through the total weight *W*_tot_, i.e.

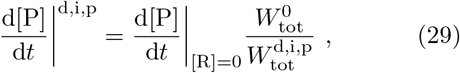

From Equations 5 and 6 we can also see that the weight is a linear function of the inhibitor concentration (see also Table 2). Again making use of the definition of the IC_50_ through Equation 9, we find

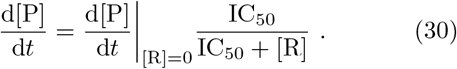

To examine this, the fluctuating inhibitor concentrations discussed in section 3.1.4 and chosen substrate concentrations of the model will be used to illustrate whether *k*_cat_, *K*_M_, catalytic efficiency, or Equations 28 and 30 can be used to inform the selection of Abl1 inhibitors based on present mutations.

## 4 Results

### 4.1 Computational results

We implemented the time-dependent differential equations as described in section 3.1.4 for the concentrations numerically using an Euler forward algorithm and therefore chose a sufficiently small time step, 1 ms.

We first consider the time-dependence of the inhibitor concentration, and then its effect on the enzyme concentrations. The inhibitor concentrations are shown in Figure 2 for the first 10 days of treatment. Each dose of the inhibitor a patient takes has a short absorption curve (increase in concentration) and a longer elimination curve (decrease in concentration). Initially, there is a build-up of concentration in the system as the absorption process is much quicker than the elimination process. After a few days, the concentration enters a periodic steady-state where the concentration fluctuates between a regular maximum and minimum with every daily dose. This is most clearly seen in the imatinib concentration in Figure 2. This variation of the inhibitor concentrations leads to variations in the concentrations of the different states of the Abl1 enzyme. Three examples of this effect can be seen in Figure 3. These examples show different responses to the addition of an inhibitor in the system. Figure 3(a) (combination of WT Abl1 and imatinib) reflects a slow, small effect of the addition of the inhibitor. In comparison, Figure 3(b) (combination of Y253H-E255V mutant Abl1 and ponatinib) shows an immediate and sustained effect to the concentrations of the different enzyme states in the system. A system that exhibits very little change after the introduction of the inhibitor can be seen in Figure 3(c), showing that dasatinib has very little influence on the states of the Abl1 which carries an T315I mutation.

**Figure 2:**
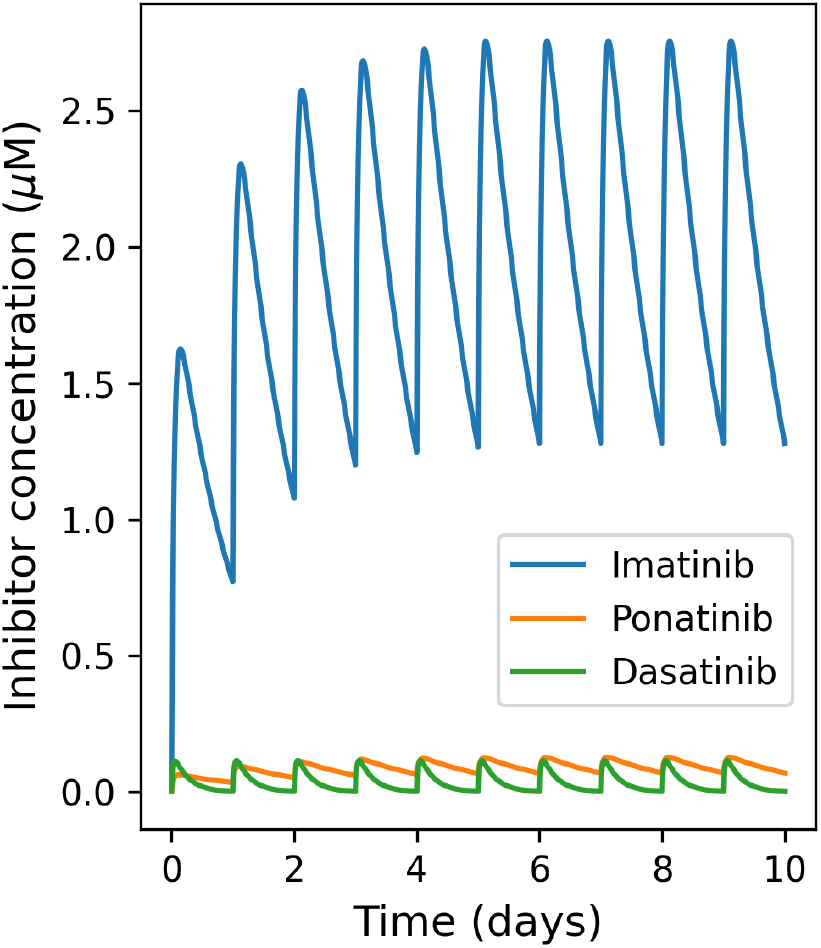
Inhibitor concentrations over the first 10 days of daily treatment doses, calculated as outline in section 3.1.4. Each inhibitor concentration increases with each daily dose (absorption), then decreases (elimination). After a build up in the system the concentration reaches a steady-state phase with a consistent minimum and maximum that the concentration fluctuates between. This is most obviously seen with imatinib, but also occurs with ponatinib and dasatinib. Dasatinib has the smallest elimination half-life, and thus reaches steady-state in the shortest time. Also observable here is that imatinib is used in much larger doses, than ponatinib and dasatinib.

**Figure 3:**
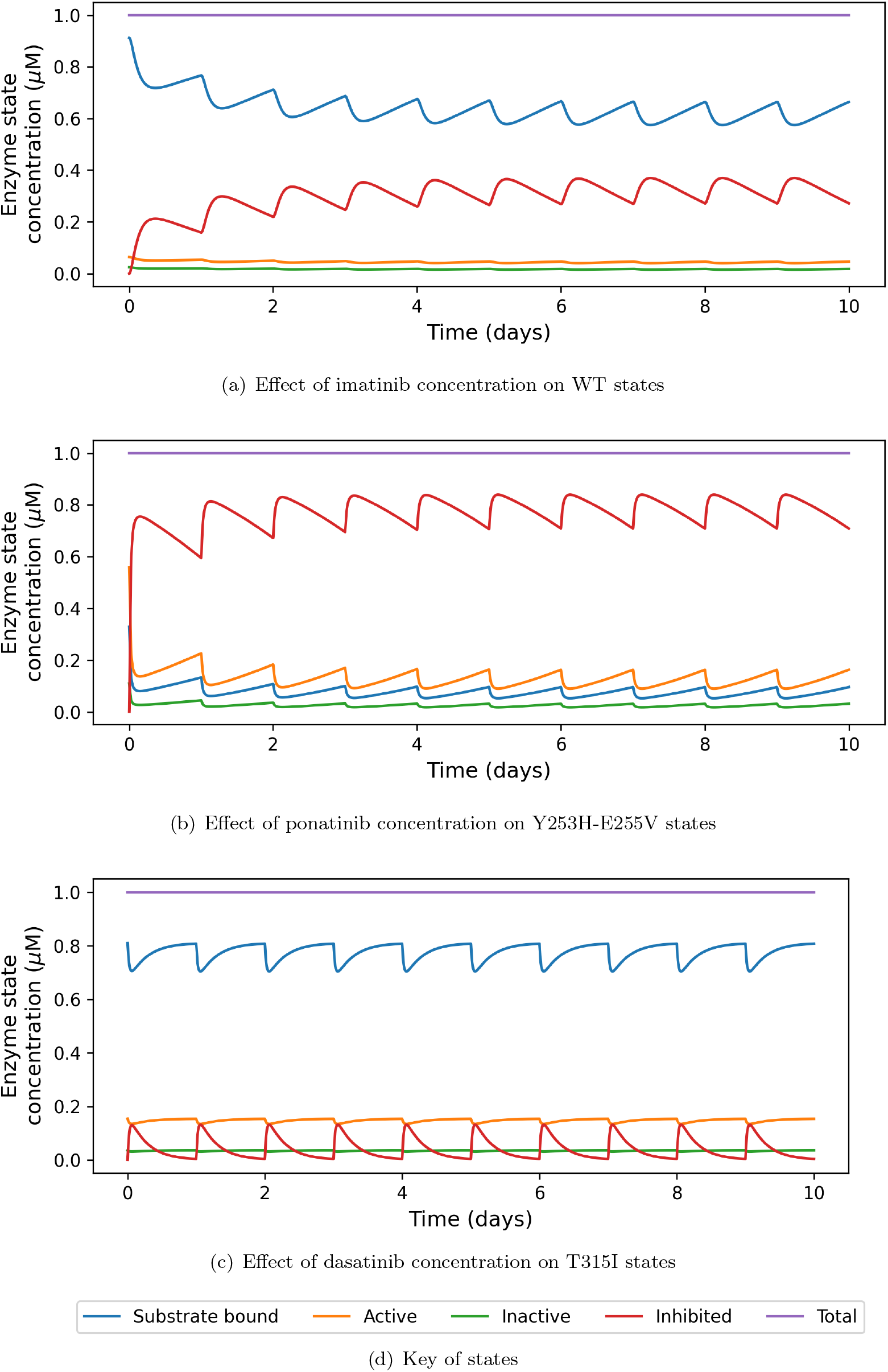
Effect of Abl1 inhibitor Concentrations on the enzyme states in the system. (a) A gradual effect as the inhibitor concentration builds up to its steady-state and the change in substrate bound enzymes from around 0.9 *µ*M to around an average of 0.6 *µ*M. (b) Here, the growth to steady-state is less visible, but the reduction in substrate bound enzymes and increase in inhibitor bound enzymes is much more dramatic. (c) Here the change in enzyme states varies, but not by a large amount, indicating that the effectiveness of dasatinib against mutation T315I is low.

As a consequence of these changes in the enzyme state concentrations, we see changes in the product formation rates, as shown in Figure 4. These changes in product formation rate do not lead to a clear explanation of the link between product formation rate and resistance to treatment. For example, in Figure 4(a), all the mutations are expected to show resistance to imatinib, but E255V shows a product formation rate lower than the WT and T315M shows similar results to the WT.

**Figure 4:**
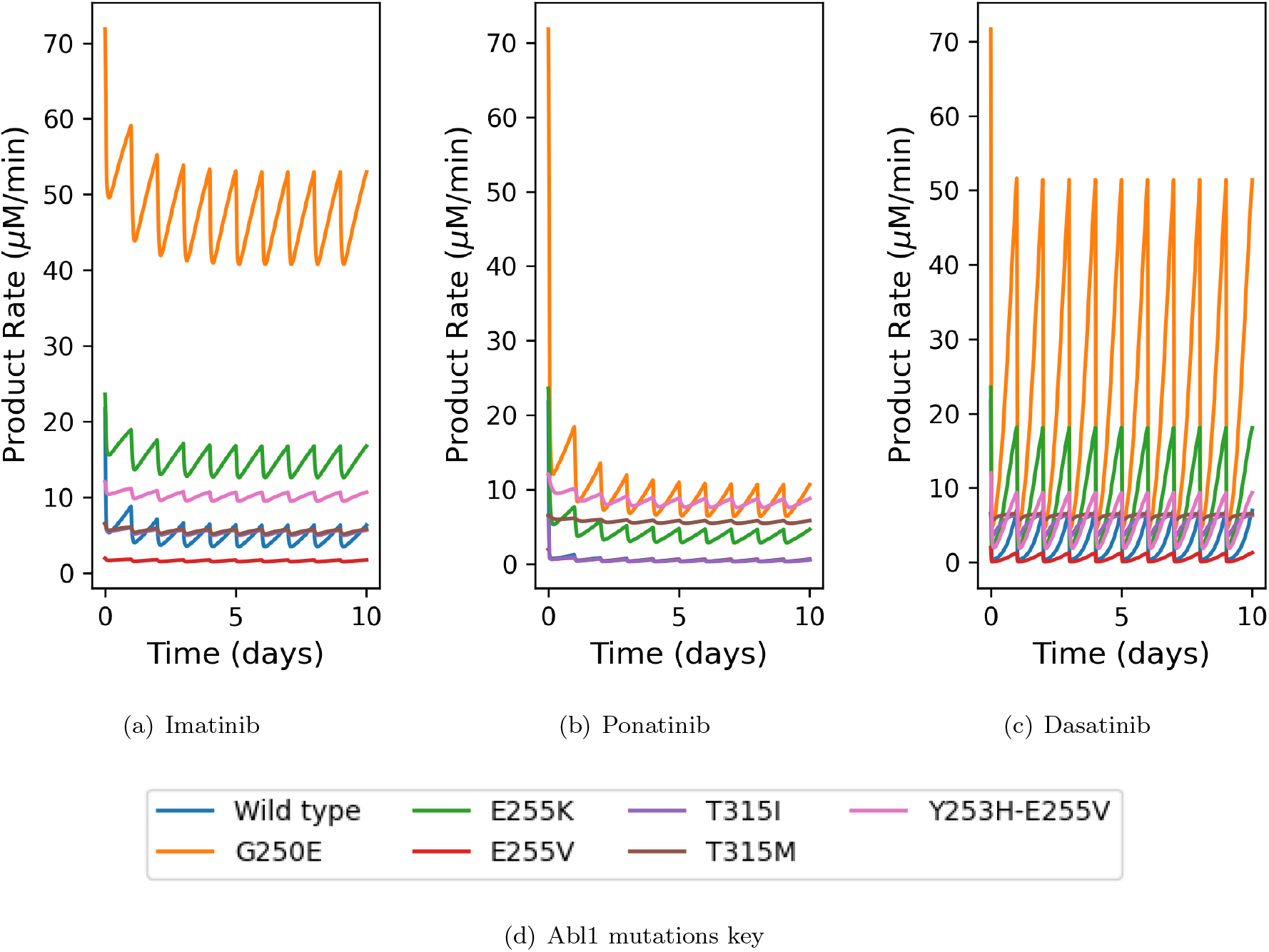
The product formation rates of the wild-type and six mutant Abl1 enzymes over ten days of initial treatment.

To further examine the cause of the root of drug resistance, we compare the “inhibitory reduction prowess” (IRP) of each mutation for each inhibitor. This is the difference in product formation rate in the steady-state halfway between two doses, compared with the initial product formation rate without inhibitor as a percentage of this initial product formation rate:

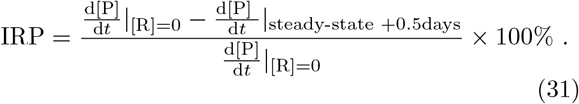

A higher value of IRP indicates that the treatment is successful at reducing the product formation rate. Since our simulations show that the steady-state is definitely reached on the 10th day, we extract the IRP from our simulations at 9.5 days since the first dose. Figure 5, shows this “inhibitory reduction prowess” and Figure 6 provides a heat map of the IRP compared with the associated resistances.

**Figure 5:**
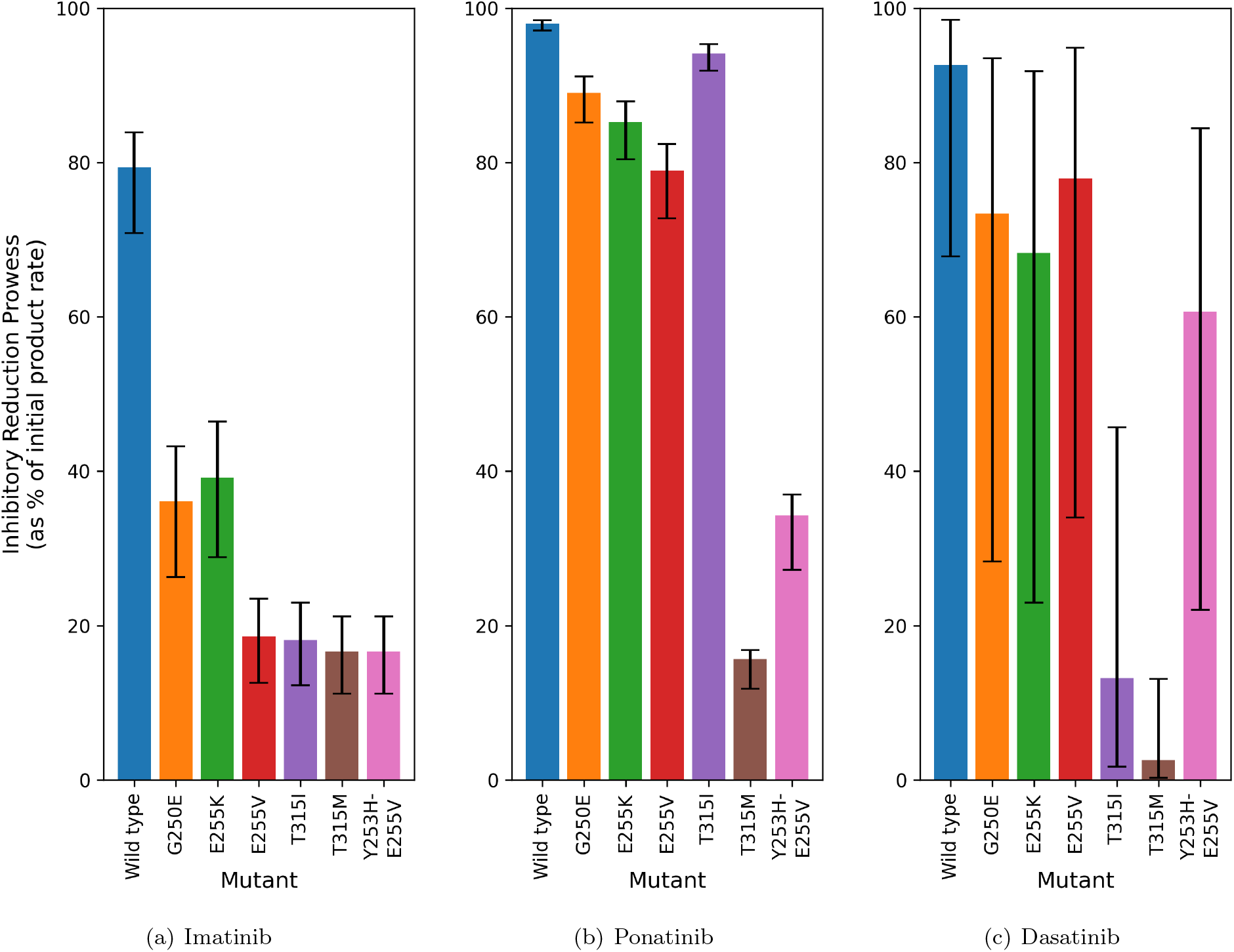
We define the term “inhibitory reduction prowess” (IRP) as the percentage by which the product formation rate is reduced by from the initial point before inhibitor is introduced to the system. A low IRP indicates a resistant mutant. The bar height is the IRP obtained from the midday level of the product formation rate on day 10 and the error bars give the range from the lowest and highest product formation rates.

**Figure 6:**
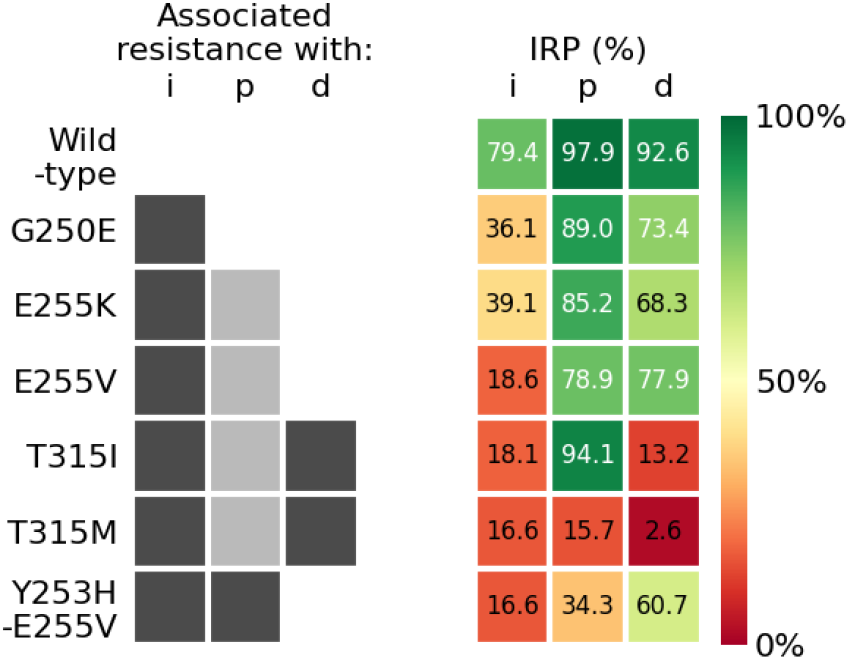
A heat map to compare the associated resistances outlined in Table 1 and [10] with the IRP values. The drugs imatinib, ponatinib, and dasatinib are represented by i, p, and d, respectively. The darker grey boxes indicate which mutations and combinations of mutations are associated with resistance to that inhibitor drug. The lighter grey boxes for ponatinib indicate that those mutations are associated with resistance when in combination with other mutations.

### 4.2 Choosing between treatments in the case of Abl1 inhibitor resistance using *k*_cat_, *K*_M_ and IC_50_ values

We now turn to the ideas discussed in section 3.2. We compare numerical evaluations of the various indicators of resistance to what is known about resistance of the various mutation/drug combinations [10].

We first consider indicators of resistance of the mutations to treatment in general that are independent of the inhibitor: the fractional part of Equation 28 ([S]*/*([S] + *K*_M_)), and the values of *k*_cat_, *K*_M_, and catalytic efficiency (*k*_cat_*/K*_M_). We also examine the information that is dependent on the inhibitors: the current method of calculating a relative 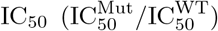 and a proposed effective IC_50_ ratio - the fractional part from Equation 30 (IC_50_*/*(IC_50_ + [R])).

Figure 7 shows information about associated resistance from Table 1 in comparison to values of the inhibitor independent information under investigation: *k*_cat_, *K*_M_, catalytic efficiency, and [S]*/*([S] + *K*_M_). The association to resistance for each mutation and drug combination is shown with grey boxes - dark grey indicates an association with resistance; light grey indicates that there is an association with this mutation when it is in combination with another mutation; and no colour when there is no resistance association. Although the values are listed, simple colour scales are added for *k*_cat_, *K*_M_, catalytic efficiency, and [S]*/*([S] + *K*_M_) to emphasise the size of the values to increase clarity of their link to associated resistance. Each colour scale shows smaller values as lighter colours and larger values as darker colours.

**Figure 7:**
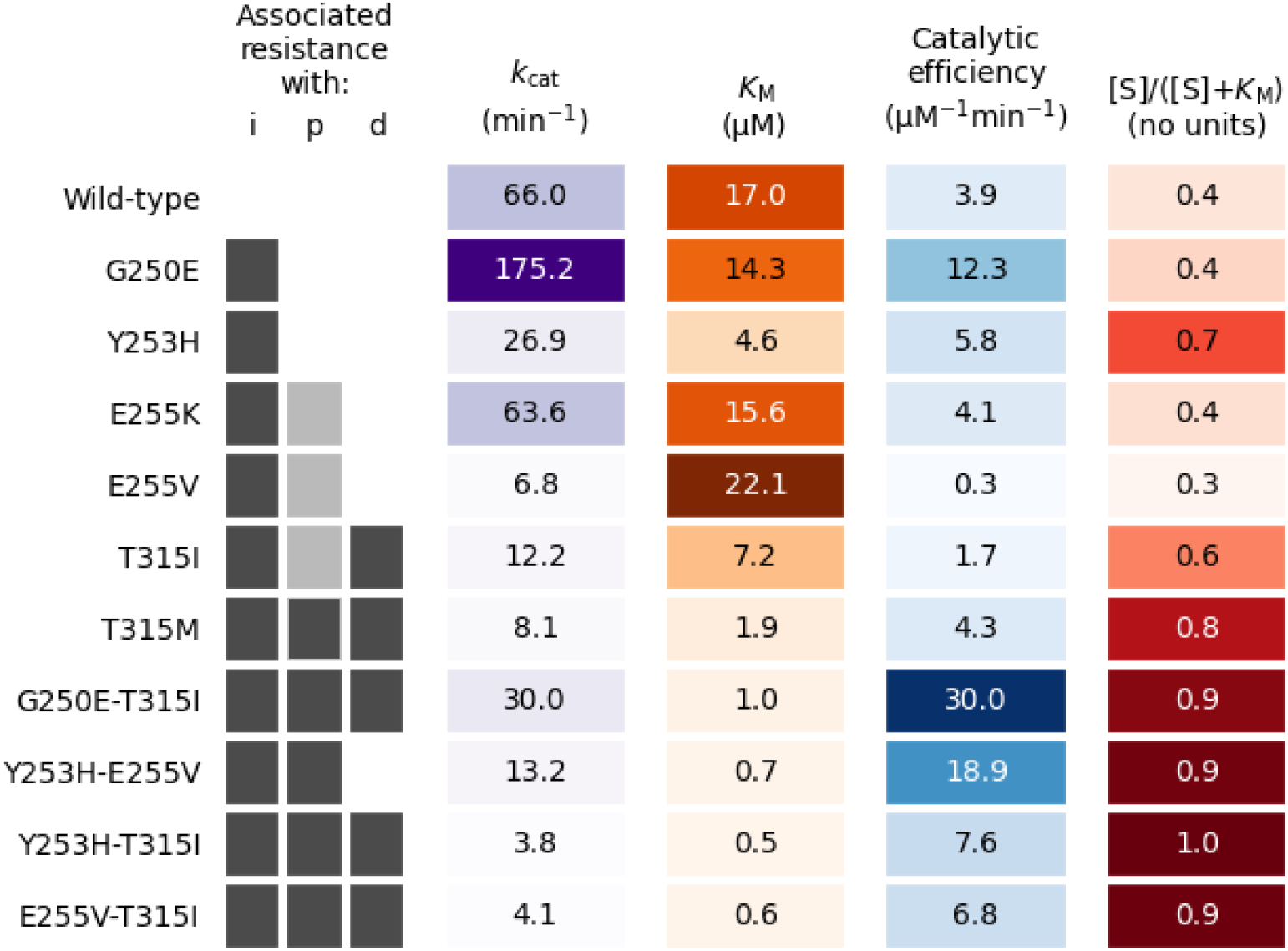
A visual comparison of the associated resistance data from Table 1 with the values of *k*_cat_, *K*_M_, catalytic efficiency (*k*_cat_*/K*_M_), and [S]*/*([S] + *K*_M_) for each mutation of Ab1. The drugs imatinib, ponatinib, and dasatinib are represented by i, p, and d, respectively. For the resistance association, the darker grey boxes indicate which mutations and combinations of mutations are associated with resistance to that inhibitor drug. The lighter grey boxes for ponatinib indicate that those mutations are associated with resistance when in combination with other mutations [10]. The colour scales for *k*_cat_, *K*_M_, catalytic efficiency, and [S]*/*([S] + *K*_M_) are simple gradient scales to emphasise the values visually with each column’s smallest value as the lightest colour and the largest value as the darkest.

As may be expected of values that have no dependence on the inhibitor type, there is no distinct cut-off on what determines resistance with the values examined in Figure 7. There are, however, some general trends that can be observed. Out of the three drugs we examine in this work, it appears that lower values of *k*_cat_ and *K*_M_ could indicate a higher chance of that mutation being resistant to a higher number of drugs.

Conversely, the opposite trend emerges for the catalytic efficiency values. This vague correlation of number of inhibitors the mutation is resistant to and size of the catalytic efficiency seems much weaker than the correlations of *k*_cat_ or *K*_M_. The values of [S]*/*([S] + *K*_M_) also show very limited correlation with higher values increasing the number of inhibitors the mutation will be associated with resistance to.

The values that are dependent on the inhibitor show much more promise. An examination of the fractional part of Equation 30 across a full day in steadystate of inhibitor concentration is shown in Figure 8. There is a clear divide between the resistant and non-resistant mutants with imatinib and ponatinib (Figures 8(a) and (b)). This divide appears to be at around IC_50_*/*(IC_50_ + [R]) = 0.5, which is where the inhibitor concentration and the IC_50_ are equal. However, the dasatinib results (Figure 8(c)) show that this divide is not strictly obeyed. Due to dasatinib’s shorter elimination half-life compared to imatinib and ponatinib, the fluctuation between the maximum and minimum points in its concentration are much larger relative to its average. The mutations that are associated with resistance to dasatinib (T315) are above this [R] = IC_50_ threshold, as seen with the other two inhibitors. What is different here is that a number of the mutations that are not associated with resistance spend a significant amount of time with IC_50_*/*(IC_50_ + [R]) *>* 0.5, because the inhibitor concentration has decreased a lot.

**Figure 8:**
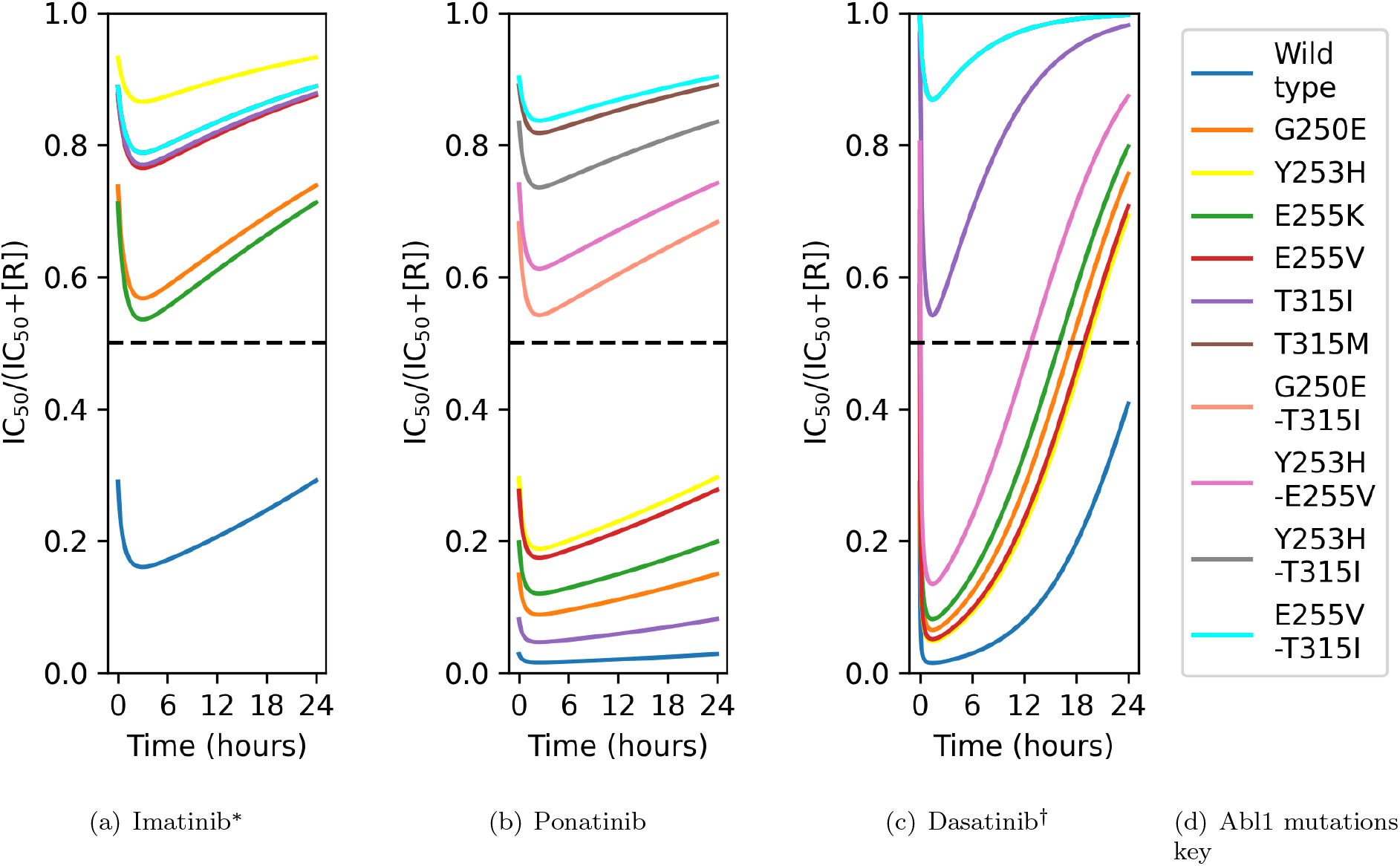
How the variation in inhibitor concentration throughout one day affects the value of IC_50_*/*(IC_50_ +[R]) for different Abl1 inhibitors. The point at which [R] = IC_50_ is marked with a dashed line. (a) Imatinib: G250E, E255K, E255V, T315I, T315M, and Y253H and their compound mutations are expected to show resistance. *T315M and all the compound mutations have the same IC_50_ value and can be represented by the result for E255V-T315I. (b) Ponatinib: Compound mutations including E255K, E255V, and T315I are expected to show resistance. (c) Dasatinib: T315I and T315M are expected to show resistance. ^†^T315M and all compound mutations containing T315I have the same IC_50_ value and can be represented by the result for E255V-T315I.

Figure 9 shows the computational results compared with current methods of choosing secondary treatment when a mutation has been reported in resistant patients. Considering the fraction 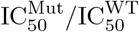, for a given inhibitor, a smaller value implies that the inhibitor is a better treatment for that mutation than for a mutation where the fraction results in a higher value. In other words, it can describe which mutations that inhibitor is suited to for treatment. It does not, however, give adequate description of which inhibitor would provide the best outcome for a given mutation. The results in Figure 9 show no clear threshold in what values of 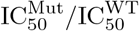 determine an outcome of resistance. Take for example the results for mutation E255K (green markers): ponatinib has the highest average IRP, but the largest value of relative IC_50_. It would appear, in terms of relative IC_50_, that dasatinib and imatinib would make better treatments for the E255K mutation; thus demonstrating how misleading these values are in comparing treatment options.

**Figure 9:**
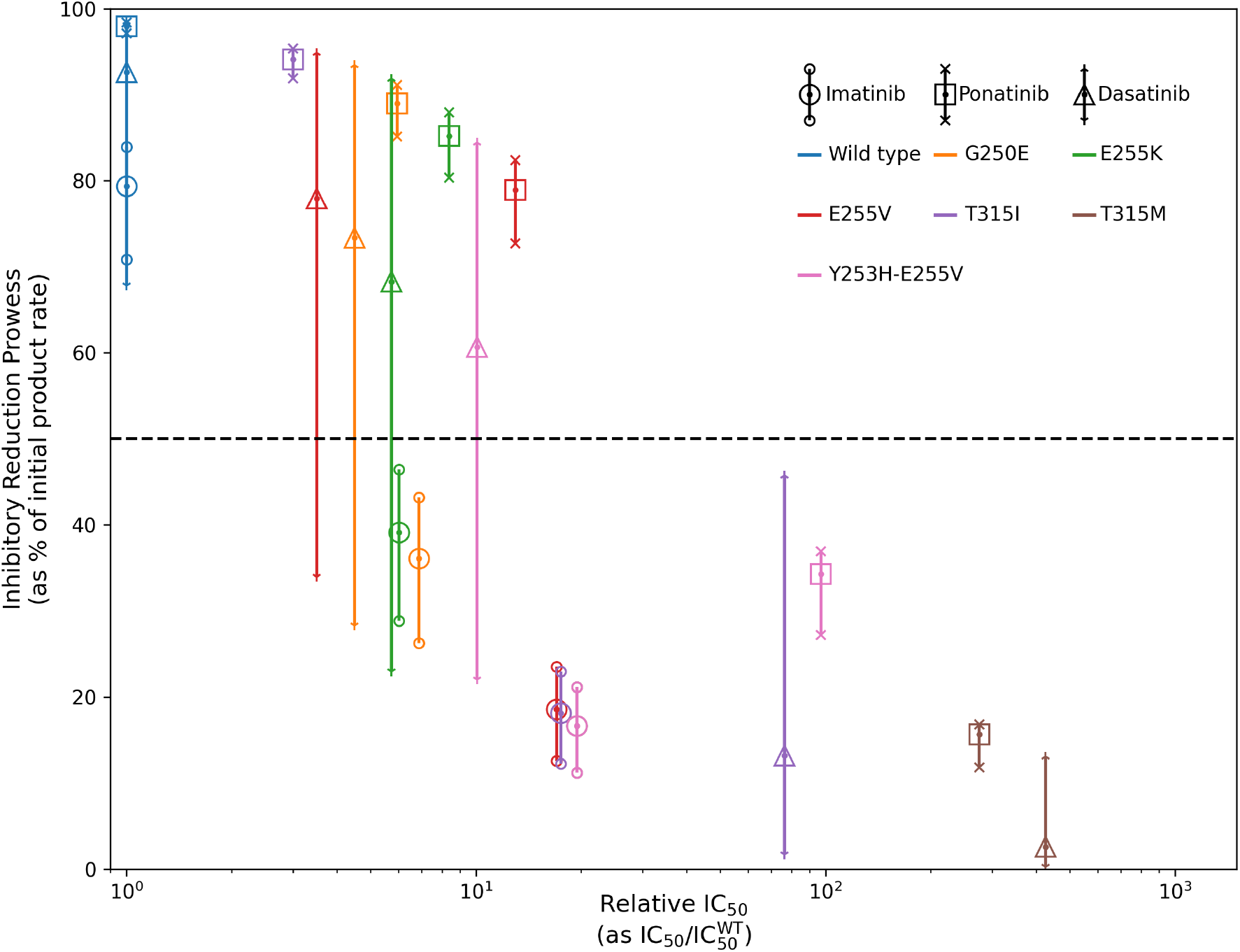
The relative IC_50_, as weighted by the IC_50_ of the WT for that Abl1 inhibitor 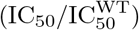 plotted against the IRP for each combination of mutation and drug. A line where IRP=50% shows a boundary, discussed previously, below which that combination of mutation and inhibitor displays resistance. The plotted points are the midday IRP on day 10 and the error bars give the range of the IRP for day 10.

Conversely, Figure 10 shows a much clearer divide in effective IC_50_ ratio and resistance. Calculating effective IC_50_ ratio as IC_50_*/*(IC_50_ + [R]) provides a way to compare between treatments for a given mutation.

**Figure 10:**
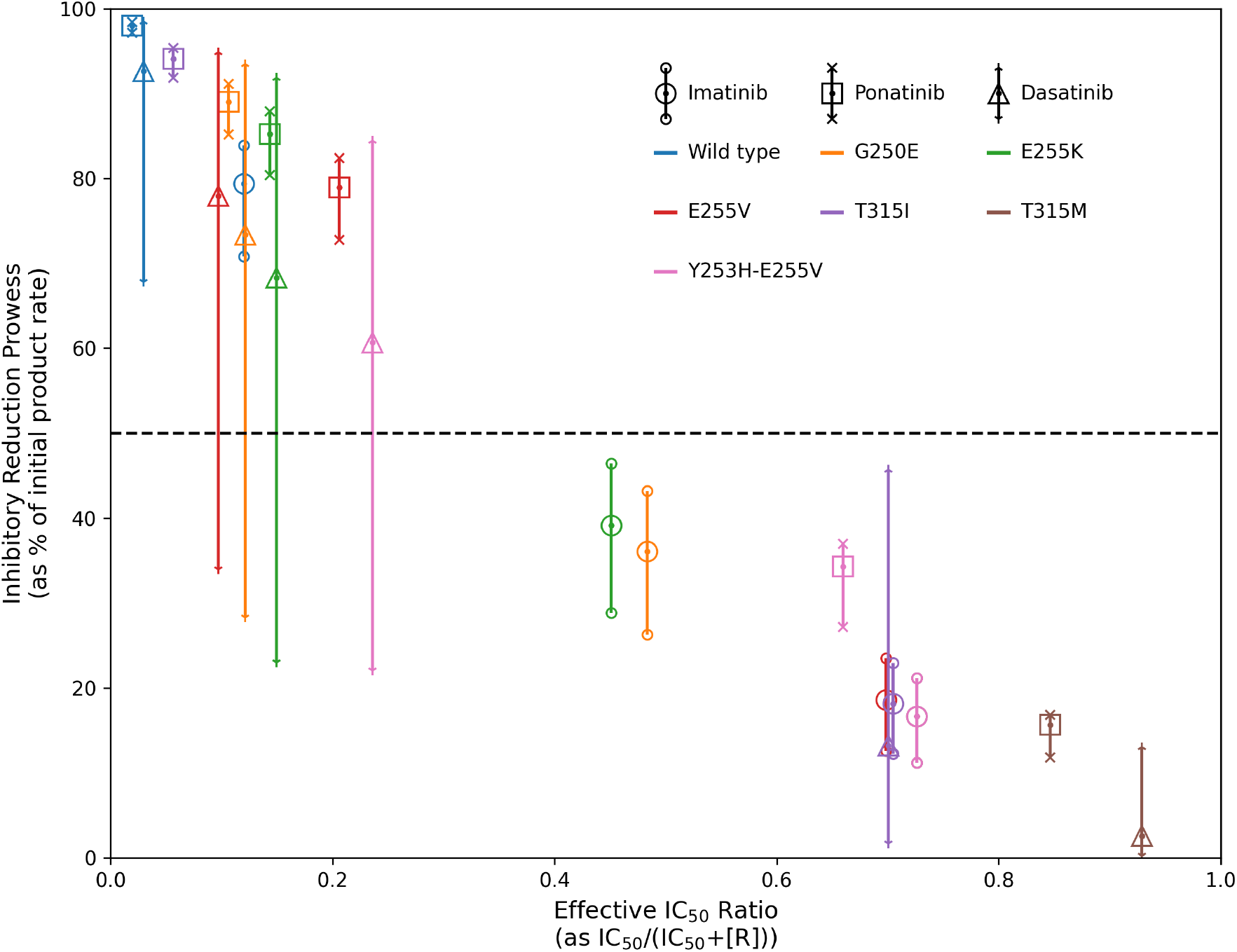
The effective IC_50_ ratio (as IC_50_*/*(IC_50_ +[R])) plotted against the IRP for each combination of mutation and drug. A line where IRP=50% shows a boundary, discussed previously, below which that combination of mutation and inhibitor displays resistance. The plotted points are the midday IRP on day 10 and the error bars give the range of the IRP for day 10.

## 5 Discussion

In this study, we have developed a model based on information of catalysis, pharmacokinetics and inhibition, in order to examine the effect of enzyme inhibitors when an enzyme develops resistance due to mutations in its sequence. The best indication of resistance is evident in the “inhibitory reduction prowess” (IRP - Equation 31), a normalised measure of the inhibitors effectiveness, rather than the actual size of the product formation rate, see Figures 4 and 5. This is most likely due to some accounting for other effects and efficiencies of the mutation in the cell. Further work for the model could include an examination of the behaviour of the model with asciminib in conjunction with each Abl1 inhibitor, as asciminib binds to a different site than the drugs examined in this work.

We also examined values and calculations that could be made to better select secondary treatment for identified mutations. As seen in Figure 7, the inhibitor independent values can help describe the chance of the mutation being resistant to an Abl1 inhibitor, but there is no clear correlation between these values and resistance. As evidenced in Figures 8 (a) and (b), resistance appears to arise when the IC_50_ value is greater than the inhibitor concentration throughout the day. This is marked in the Figures as IC_50_*/*(IC_50_ + [R]) = 0.5. Due to the short elimination half-life of dasatinib, the steady-state of the plasma concentration has a range that begins closer to zero than other Abl1 inhibitors. This allows for a relatively larger range of values for IC_50_*/*(IC_50_ + [R]), meaning that for a portion of each day, the system is in a”state of resistance” even for otherwise non-resistant mutants. This indicates that values of the average plasma concentrations of Abl1 inhibitors in patients combined with IC_50_ information could inform oncologists when selecting which second or third generation treatment to switch to once resistance arises and the mutation has been identified. Further comparison between Figures 9 and 10 shows that not only is there a clearer numerical divide in resistance for IC_50_*/*(IC_50_ + [R]) over the current method of 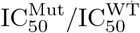, the effective IC_50_ ratio occurs over a smaller, and more specific, range and has a more linear correlation to IRP. Hence, the numerical values from this effective IC_50_ ratio could provide a clearer and more intuitive process for selecting secondary treatments. It would be possible to create a tool that could provide this information to oncologists and could have a great impact on patient outcomes.

